# Broad-scale spatial patterns of canopy cover and pond morphology affect the structure of a Neotropical amphibian metacommunity

**DOI:** 10.1101/001255

**Authors:** Diogo B. Provete, Thiago Gonçalves-Souza, Michel V. Garey, Itamar A. Martins, Denise de C. Rossa-Feres

## Abstract

Spatial and environmental processes influence species composition at distinct scales. Previous studies suggested that the landscape-scale distribution of larval anurans is influenced by environmental gradients related to adult breeding site selection, such as pond canopy cover, but not water chemistry. However, the combined effects of spatial, pond morphology, and water chemistry variables on metacommunity structure of larval anurans have not been analyzed. We used a partial redundancy analysis with variation partitioning to analyze the relative influence of pond morphology (e.g., depth, area, and aquatic vegetation), water chemistry, and spatial variables on a tadpole metacommunity from southeastern Brazil. We predict that the metacommunity will be spatially structured at broad spatial scales, while environmental variables, mainly related to adult habitat selection, would play a larger role at fine spatial scales. We found that broad-scale spatial patterns of pond canopy cover and pond morphology strongly influenced metacommunity structure. Additionally, species composition was spatially autocorrelated at short distances. We suggest that the reproductive behavior of adult anurans is driving tadpole metacommunity dynamics, since pond morphology, but not water chemistry affects breeding site selection by adults. Our results contribute to the understanding of amphibian species diversity in tropical environments.

## Introduction

A main goal of ecology is to find causal explanations for patterns of distribution and relative abundance of species (Vellend 2010). Traditionally, environmental gradients have been deemed as major determinants of species distribution. However, spatial gradients may also influence the assembly of ecological communities, for example as surrogates for dispersal ability (Beisner et al., 2006; McIntire & Fajardo, 2009; Landeiro et al., 2011). Accordingly, spatial dynamics have continuously been incorporated into community ecology through several frameworks (*e.g.*, Leibold et al., 2004). The primary framework is metacommunity theory, which has advanced our understanding of how dispersal and patch heterogeneity influence species abundance and distribution in a set of local communities (Leibold et al., 2004). In addition, the metacommunity concept represents a multiscale approach to test the joint effects of environmental and spatial processes on species diversity.

Recent studies (Logue et al., 2011; Winegardner et al., 2012) have claimed for an integrated view of the metacommunity perspectives (Leibold et al., 2004). The assumptions and predictions of these perspectives differ depending on the amount of dispersal rate and patch heterogeneity at play in the metacommunity. For instance, the mass effect perspective predicts that species composition varies with both space and environment, when environment is spatially structured (Chase et al., 2005). However, species sorting also modulates organismal responses to environmental heterogeneity (Leibold et al., 2004), especially when organisms track environmental characteristics related to their fitness, such as breeding site selection (Resetarits et al., 2005). Nonetheless, data from a number of empirical studies have conformed usually to more than one perspective (Cottenie, 2005; Logue et al., 2011).

Spatial patterns in metacommunities may arise from neutral- and niche-based mechanisms. Spatial distribution of species under niche-based mechanisms is driven by their distinct demographic characteristics as a result of differential responses to the environment (stabilizing mechanisms; Chesson, 2000). According to this view, individuals occupy sites along environmental gradients in order to maximize their fitness. Conversely, under neutral dynamics species are ecologically equivalent (equalizing mechanisms) and have the same chance to give birth, die, migrate, and speciate (equalizing mechanisms; Chesson, 2000), and thus their abundance is dictated solely by stochastic demographic fluctuations. Consequently, species distribution would be random, but spatially autocorrelated, due to dispersal limitation of organisms (equalizing mechanisms; Leibold et al., 2004). Nonetheless, discerning the scale at which spatial processes influence ecological patterns remains a challenge for ecologists (Levin, 1992; Landeiro et al., 2011).

The joint influence of environmental and spatial processes on metacommunity structure has been largely studied on theoretical grounds (Leibold & McPeek, 2006). As a result, empirical studies on the effects of environmental and spatial processes on metacommunity structure of freshwater organisms are scarce, especially in tropical environments (Logue et al., 2011). For instance, recent experiments with aquatic microorganisms found interacting effects of local environmental factors and dispersal on species composition and diversity (Cadotte, 2006; Altermatt et al., 2011). Pond organisms with complex life cycles, such as anuran amphibians, are a useful model system to test metacommunity theory, since they are subjected to different spatial and environmental processes at the adult and larval stages. Most amphibians are dispersal-limited organisms that require wet environments to live and reproduce (reviewed in Smith & Green, 2005; Wells, 2007). Additionally, several anuran amphibians exhibit a marked philopatric behavior, breeding in the same pond in which they emerged as juveniles (Semlitsch, 2008), which reinforces the potential effects of both environmental and spatial processes on anuran metacommunity structure. This complex habitat selection behavior coupled with specific environmental requirements may act as an endogenous ecological process (McIntire & Fajardo, 2009; Legendre & Legendre, 2012), creating spatial autocorrelation in species composition. Moreover, adults are the life stage that disperse and effectively connect breeding sites by oviposition (Wells, 2007; Semlitsch, 2008). As a result, both the spatial arrangement of water bodies and their surroundings may play a role in the distribution of pond-breeding anurans and their larvae.

The dynamics of the terrestrial ecosystem surrounding ponds strongly also affects freshwater metacommunities. One aspect of the terrestrial ecosystem that strongly influences the aquatic ecosystem is pond canopy cover, which is a key gradient affecting the distribution of several freshwater organisms, from odonate larvae (McCauley et al., 2008), to tadpoles (Skelly et al., 1999; Werner et al., 2007), to snails (Hovermann et al., 2011). Canopy cover alters light availability and leaf litter input into ponds, and consequently primary productivity and decomposition (Werner et al., 2007). As a result, larval fitness is lower in less productive, closed-canopy ponds (Schiesari, 2006). Besides primary productivity, canopy cover may affect other water chemistry variables that influence larval development, such as temperature, conductivity, and dissolved oxygen (Werner & Glennenmeier, 1992). Consequently, few species tolerate and colonize closed-canopy ponds, but those that do benefit from lower exploitative competition (Schiesari, 2006).

On the other hand, the influence of water chemistry on freshwater communities has been rarely investigated at the metacommunity scale (*e.g.*, Hecnar & M’Closkey, 1996; Hajek et al., 2011). Several water chemistry variables modulate tadpole survival, development, and time to metamorphosis. Conductivity is often correlated with dissolved ions and solids, mineral particles, disturbance or eutrophication that affect osmoregulation and larval development (Welch & MacMahon, 2005). Low pH disrupts the Ca:Na ratio, directly influencing survival. Temperature and dissolved oxygen affect metabolic rates and directly influence time to metamorphosis and larval fitness (reviewed in Ultsch et al., 1999). Turbidity is related to the trophic status of ponds (Scheffer et al., 1993). Hence, water chemistry may represent the influence of local environment on tadpole relative abundance, as expected under the species sorting perspective.

Taken together, the aforementioned environmental variables seem to differentially affect adult anurans and their larvae. On the one hand, the distribution of adult anurans seems to be more influenced by spatial processes and landscape attributes of the terrestrial ecosystem (Resetarits et al., 2005; Binckley & Resetarits, 2007; Wells 2007). On the other hand, larval development, growth, and abundance might be directly influenced by local water chemistry (Ultsch et al., 1999), although any metacommunity-scale process on adults will probably affect their larvae. However, to the best of our knowledge, empirical studies have not tested the combined effects of spatial structure, water chemistry, and pond morphology on amphibian species composition, especially in diverse, tropical environments (see also Logue et al., 2011). In particular, the complex life cycle characteristic of amphibians has the potential to simultaneously test whether different perspectives in metacommunity theory could explain community assembly.

Therefore, we hypothesized that these two sets of environmental variables represent alternative mechanisms driving metacommunity dynamics. Following McIntire & Fajardo (2009), we argued that pond morphology and canopy cover could be proxies for the effect of adult breeding site selection at the metacommunity scale, while water chemistry represented the local effects of pond environment, by influencing tadpole development and survival. In this framework, closed-canopy ponds are harsh environments, acting as a sink, whereas open-canopy ponds are more favorable, source environments. Therefore, we hypothesize that a mass effect may be involved in maintaining populations in closed-canopy ponds, which disperse from open-canopy. Concurrently, species sorting may also play a role in amphibian metacommunities, given the low dispersal ability of adults, and their tendency to track environmental characteristics related to breeding sites. As a result, we expect that pond morphology and canopy cover influence species composition at broad scales, while water chemistry would be more significant at fine spatial scales.

## Material and methods

### Study area and sampling

This study was carried out in the Serra da Bocaina National Park (22° 40’ to 23° 20’ S; 44° 24’ to 44° 54’ W), at the border between the states of São Paulo and Rio de Janeiro, southeastern Brazil (Fig. 1). Our samples were collected on the Bocaina highlands (1,500 m a.s.l.), in São José do Barreiro, São Paulo, and extended over an area of approximately 11 Km^2^ (ponds were 2.15 Km apart in mean, maximum 6 Km). The climate in this region is of the type Cwb (humid subtropical highland) according to the Köppen-Geiger classification (Peel et al., 2007), with moderated temperatures, dry winters (between April and September) and warm summers (from October to March). The annual rainfall varies between 400 and 2100 mm, with mean annual temperature of 22° C. The breeding season for the majority of adult anurans is during the rainy season (M.V. Garey, unpub. data). Thus, we sampled ponds in this period in order to maximize the chance of collecting most of species.

**Fig. 1.**
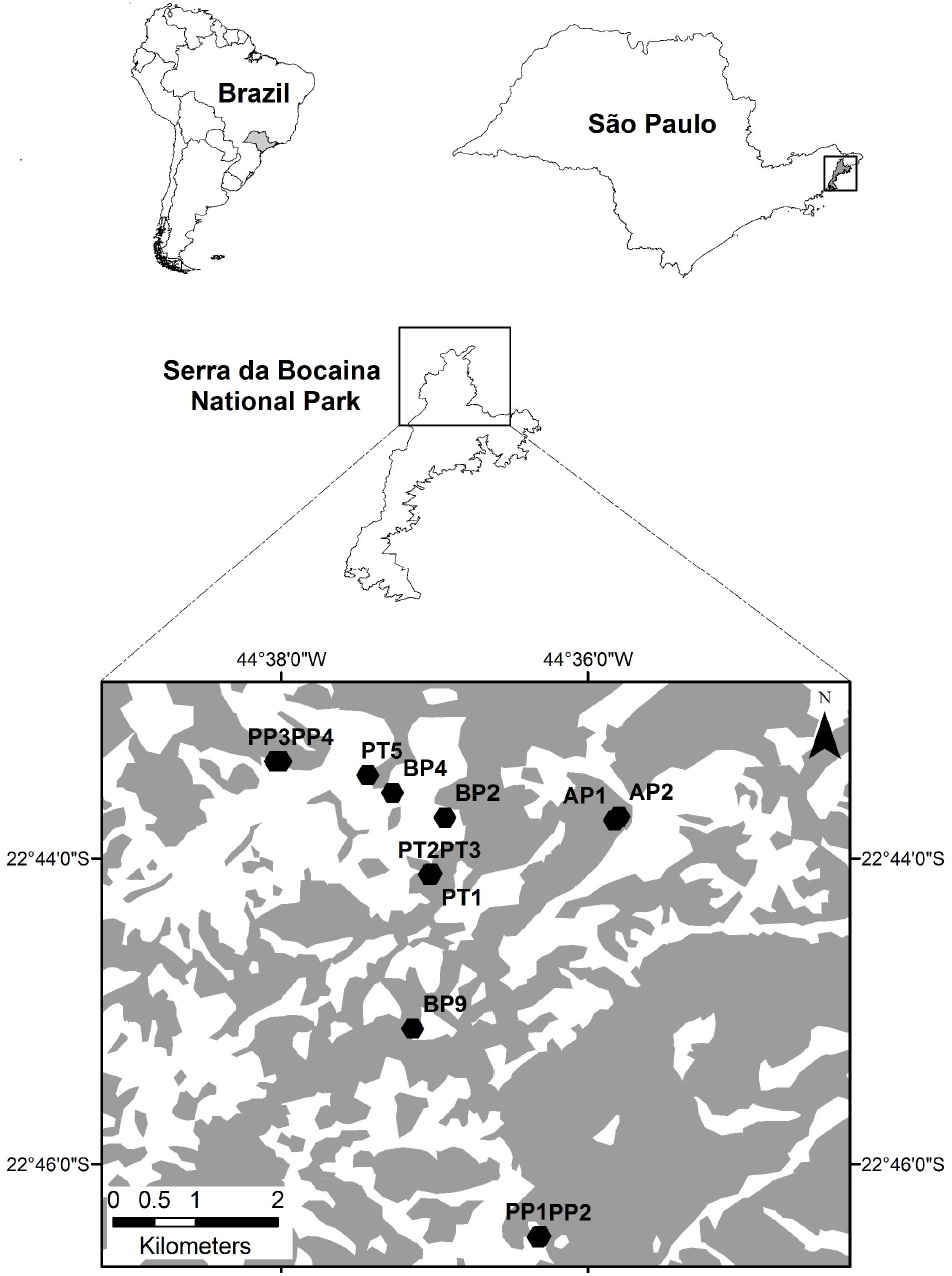
Map showing the location of the study area and the spatial distribution of sampled ponds in the Serra da Bocaina National Park, southeastern Brazil. Gray areas represent primary or secondary forest; white areas represent open areas.

We sampled tadpoles using a hand dipnet in 13 water bodies with different morphologies (e.g., area, depth, and aquatic vegetation) and varying degrees of canopy cover, monthly between August 2008 and January 2009. All ponds were sampled within a one-week period each month. All ponds were fishless, and 3 out of 13 held water during only three months, all others were permanent ponds. Blind sweeps were made along the entire margins of water bodies, with effort proportional to surface area (Skelly & Richardson, 2010). Tadpoles were fixed in the field with 10 % formalin. In addition, adult anurans were acoustically monitored monthly to record the species present in each water body, which would help in tadpole identification.

Prior to tadpole sampling, we recorded two sets of pond characteristics: (i) water chemistry variables, namely: conductivity (mS/cm), dissolved oxygen (mg/l), pH, temperature (°C), and turbidity (NTU) using a Horiba U-10 multiparameter water quality checker; (ii) pond morphology variables: surface area (m^2^), maximum depth (m), and aquatic vegetation (%). We estimated aquatic vegetation visually by dividing pond surface into quadrants; the percentage of aquatic vegetation was used as a proxy for habitat structural complexity, varying from 0 (low complexity) to 100 % (high complexity). We also recorded pond canopy cover (%) using a spherical densitometer (Forestry Suppliers, Jackson, MS, U.S.A.); measurements were taken in four directions (N, S, E, W), and the center of the pond. Pond morphology variables were measured only once at the peak of the rainy season, whereas water chemistry was measured during all sampling period. Prior to analyses, we standardized environmental variables to zero mean and unit variance to account for their different scales of measurement. Calculations were implemented in *vegan* package (Oksanen et al., 2011) in R version 2.13.2 (R Core Team, 2011).

### Spatial variables

We used distance-based Moran’s Eigenvector Maps (dbMEM; Dray et al., 2012; Legendre & Legendre, 2012) to describe spatial structures. We computed dbMEM eigenvectors from a truncated geodesic distance matrix obtained with R package *fields* (Furrer et al., 2011). The longest distance (2.887 Km) connecting two ponds in a minimum spanning tree of sampling sites (Fig. S1) was used as a threshold to truncate the distance matrix. This procedure produced 7 dbMEMs. Analysis was conducted using R package *PCNM* (Legendre et al., 2010). By concentrating most of the variation, the first eigenvectors usually describe broad spatial structures, while last eigenvectors describe fine spatial structures (Dray et al., 2012; Legendre & Legendre, 2012). Further, we implemented a forward selection procedure with double-stopping criteria (Legendre & Legendre, 2012) in R package *packfor* (Dray, 2009) to only select dbMEMs that significantly explained the variance in the species composition matrix. This procedure recovered 5 dbMEM (accumulated R^2^_adj_ = 0.414; Fig. S2-S3). dbMEM 1 and 2 were arbitrarily classified as broad-scale variables, dbMEM 4 as medium scale, and dbMEM 5 and 7 as fine-scale variables (Fig. S2). The first two dbMEMs modeled positive spatial autocorrelation (Moran’s *I* = 0.291 and 0.084, respectively), while the last three modeled negative autocorrelation (Moran’s *I* = -0.218, -0.068, and -0.033, respectively). All eigenvectors were used as spatial variables in the following analysis, since excluding those with negative Moran’s *I* seems not to change overall results (see Legendre & Legendre, 2012).

### Data analyses

For all subsequent analyses, we transformed the total counts of species using the Hellinger transformation (Legendre & Legendre, 2012) to homogenize variation among species abundances. None environmental variable had a Variation Inflation Factor higher than 3. Thus, all were included in the further analysis. We employed a variation partitioning approach to disentangle species response to environmental (pond morphology, canopy cover, and water chemistry) and spatial variables (dbMEMs) driving metacommunity structure (Cottenie, 2005; Legendre & Legendre, 2012). Despite some recent criticism (Gilbert & Bennett, 2010; Smith & Lundholm, 2010; but see Diniz-Filho et al., 2012), this analytical technique remains useful to separate variation in species composition generated by environmental gradients, spatial autocorrelation, and spatially structured environmental gradients (shared variance). We determined the pure and shared effects of water chemistry, pond morphology, and pond spatial network (dbMEM) on species composition using a partial Redundancy Analysis (pRDA; Legendre & Legendre, 2012), with adjusted canonical R^2^ values (Legendre & Legendre, 2012).

Since we sampled ponds repeatedly, we used each pond and month as factors in the pRDA and then performed a randomization stratified within ponds (Lepš & Šmilauer, 2003), with 999 Monte Carlo randomizations to determine significance. We also built a Mantel correlogram to investigate the spatial pattern of species composition and also correlograms to analyze the spatial patterns of environmental variables. Analyses were implemented in R packages *vegan* (Oksanen et al., 2011) and *spdep* (Bivand, 2012). To identify the environmental variables related to each spatial model, we regressed each dbMEMs (explanatory variables) against pond morphology and water chemistry (predictor variables; Borcard et al., 2004). Linearity was checked visually.

## Results

We recorded tadpoles of 15 amphibian species belonging to 5 families. The most abundant species were *Rhinella icterica* (3,532 individuals), *Hypsiboas* sp. (aff. *polytaenius*) with 2,339 individuals, and *Scinax* sp. (aff. *hayii*) with 2,318 individuals. The mean species richness per pond was 2.8 (± 1.45 SD; range = 2 - 9). *Chiasmocleis mantiqueira* and *Scinax* sp. (aff. *obtriangulatus*) were represented by only two individuals and were not included in further analysis, since they could affect pRDA.

Both environmental variables and spatial distribution of ponds affected metacommunity structure. Pond morphology and canopy cover accounted for 16 % (*P* = 0.005), whereas water chemistry did not explain a significant proportion of the variation in species composition (R^2^_adj_ = 0.00; *P* = 0.75; Fig. 2). Pond canopy cover and aquatic vegetation were the two variables that mainly influenced species composition (Table 1; Fig. 3). The pure spatial component explained about 20 % (*P* = 0.005) of the variation in species composition. Residual variation represented 40 %. Furthermore, it is noteworthy that all variation (16 %) in pond morphology and canopy cover were spatially structured (Fig. 2).

**Fig. 2.**
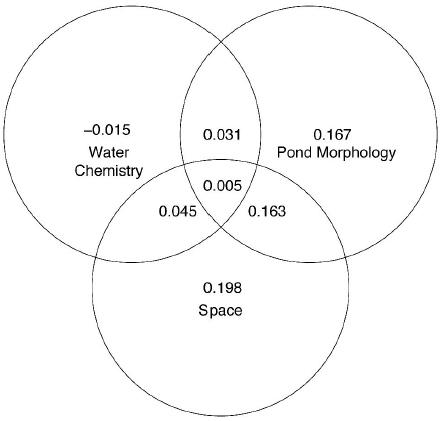
Result of variation partition with partial Redundancy Analysis showing the pure and shared influences of spatial variables, water chemistry, and pond morphology on tadpole species composition. Numbers represent adjusted canonical R^2^ values.

**Fig. 3.**
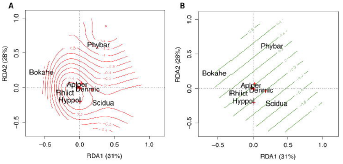
Biplot of the Redundancy analysis showing the major influence of A) pond canopy cover and B) aquatic vegetation on Hellinger-transformed data of tadpole species abundance. Isolines represent the fited smooth surface for each environmental variable, obtained with function *ordisurf* in R package *vegan*. Only species with high scores are shown. Aplper=*Aplastodiscus perviridis*, Bokahe=*Bokermannohyla ahenea*, Denmic= *Dendropsophus microps*, Hyppol=*Hypsiboas* sp. (aff. *polytaenius*), Phybar=*Physalaemus barrioi*, Rhiict=*Rhinella icterica*, Scidua=*Scinax* sp. (aff. *duartei*).

Species composition showed significant positive spatial autocorrelation in the first two distance classes (between 0.01 and 1.63 Km) and negative autocorrelation in the third and fourth distance classes (between 2.43 and 3.24 Km; Fig. S4). Ponds separated by 2.43 Km (upper limit of the third distance class) differ in composition, probably due to contrasting values of canopy cover, area, depth, and aquatic vegetation at short distances (Fig. S5). Water depth was the most important variable, explaining all spatial variables. Other variables were related to one or two broad-scale dbMEMs. Temperature, pH, and turbidity were not significantly related to any spatial variable (Table 2). Thus, the strong induced spatial dependence of pond morphology and canopy cover (R^2^_adj_ = 0.16; Fig. 2), but not water chemistry (R^2^_adj_ = 0.04; Fig. 2) are directly influencing the spatial dynamics of amphibian species. Taken together, these results suggest that species sorting and mass effects related to adult breeding site selection are shaping metacommunity structure.

## Discussion

We found that both the pure spatial and environmental component related to pond morphology and canopy cover accounted for large and significant fractions of the variation in species composition. More importantly, the gradients of pond canopy cover and morphology were spatially structured. These results point to a combination of different metacommunity processes, in which both mass effects and species sorting are playing key roles in metacommunity dynamics. Additionally, the induced spatial dependence of variables from the surrounding terrestrial ecosystem (i.e., pond morphology and canopy cover), which we hypothesized to represent the effects of adult breeding site selection, suggest that spatial dynamics of anuran larvae is essentially determined by behavioral decisions of adults in response to spatially structured environmental gradients.

All of the variation in species composition accounted for by pond morphology and canopy cover was spatially structured. We hypothesized that canopy cover and other pond morphology variables represented the effects of the surrounding terrestrial landscape on adult habitat selection (Resetarits et al., 2005; Binckley & Resetarits, 2007). In this sense, tadpole metacommunity structure seems to be essentially determined by environmental characteristics that vary at the metacommunity scale (i.e., pond morphology), in contrast to local, pond water chemistry variables. Furthermore, we also showed that the pure spatial processes (∼ 20 %) seem to play a larger role in shaping species composition, compared to pond morphology variables alone (16 %). The high variation accounted for by the pure spatial component can be attributed either to unmeasured environmental variables or pure spatial processes, such as dispersal limitation (Landeiro et al., 2011; Legendre and Legendre, 2012). Indeed, several niche-based processes predict dispersal limitation, especially when associated with spatially heterogeneous environments, demographic stochasticity, or strong, asymmetric competition (Chesson, 2000; Chase et al., 2005).

The broad-scale spatial pattern of pond morphology variables and canopy cover seems to influence tadpole metacommunity dynamics. These variables are also known to influence breeding site selection by adults (Resetarits et al., 2005; Binckley & Resetarits, 2007; Skelly et al., 1999; Werner & Glennemeier, 1999). An experimental study (Binckley & Resetarits, 2007) showed that treefrogs actively selected open-canopy ponds and deposited significantly more eggs in these ponds than in closed-canopy ponds. Additionally, treefrogs and aquatic beetles discriminated between fish and fishless ponds for oviposition sites. In our study area, canopy cover was lower in the northernmost ponds (Table 2). As a result, species inhabiting open-canopy ponds are more abundant in that region (except *D. minutus* and *B. ahenea*, which occur more widely, but are less abundant elsewhere). These results support the idea that stabilizing niche differences (Chesson, 2000) related to different environmental requirements for adult reproduction as key mechanisms structuring this metacommunity.

Contrary to our initial prediction about within-pond community structure, water chemistry variables do not seem to influence tadpole species composition. This result is in agreement with previous studies on amphibians (Hecnar & M’Closkey, 1996; Brodman et al., 2003). Other abiotic (e.g., pond canopy cover, productivity, hydroperiod) and biotic variables (e.g., presence of fish) may be more important than water chemistry in determining pond colonization by adults and survival of tadpoles. Additionally, water chemistry variables seemed not to vary in a wide range at the landscape-scale in our study area. Phenotypic plasticity mediates species coexistence in ecological communities and also their response to environmental variation (Miner et al., 2005). The high adaptive plasticity of tadpoles in response not only to predators (Relyea and Werner, 2000), but also to local environmental variables, food availability, and pond drying (Newman, 1992) is a possible mechanism responsible for the lack of influence of water chemistry on species composition.

The spatial autocorrelation in species composition at short distances seems to be driven by the induced spatial dependence of pond morphology variables, such as aquatic vegetation, depth, area, and canopy cover. However, it is difficult to tease apart if species composition is spatially autocorrelated due to dispersal limitation or due to species response to spatially structured environmental variables, or even a combination of both (Shurin et al., 2009). Dispersal ability varies among species in a metacommunity and populations of some species may be more isolated than others. Dispersal in adults is usually restricted to the surroundings of one breeding site or a cluster of ponds nearby (Semlitsch, 2008). As a consequence, it is still difficult to assign a single mechanism (e.g., dispersal limitation, philopatry) to the pure spatial component. Despite that limitation, previous studies using microsatellite markers as an indirect way to estimate dispersal in anurans (Newman & Squire, 2001 and references therein) did not find population differentiation at the scale of 2 km. These data agree with our results about spatial autocorrelation in species composition, since we also did not find a differentiation in species composition of ponds separated by 2.43 Km. Thus, despite the lack of data for dispersal ability, these results could suggest that both dispersal limitation and responses to spatially structured environmental variables can be involved in the dynamics of this larval metacommunity.

In summary, we have shown that the induced spatial dependence of pond morphology and canopy cover strongly influenced the tadpole metacommunity structure. These findings posit strong role for adult behavior in determining larval ecology. It might be possible to extrapolate our results to explain the distribution of other organisms with similar complex life cycles that represent an important link between aquatic and terrestrial ecosystems and play a key role in biomass export (Earl et al., 2011). The effect of the surrounding terrestrial landscape in determining species assembly in the adjacent, aquatic ecosystem deserves further investigation in other freshwater organisms. For example, understanding how the spatial dynamics of terrestrial landscape and species composition modulate aquatic communities may contribute to studies modeling the flux of species and subsides, trophic interactions, and metaecosystem dynamics (Massol et al., 2011). Future studies should also integrate the effects of broad- and fine-spatial scales on amphibian species with different dispersal abilities. In addition, metacommunity simulations in an environmentally homogeneous domain could evaluate how philopatric behavior and neutral dynamics create spatial patterns.

## Acknowledgments

We are indebted to M. Almeida Neto, F. Altermatt, L. M. Bini, J. W. Fox, V. L. Landeiro, G. F. Livingston, T. F. Rangel, and T. Siqueira for providing very useful insights to this manuscript. A. Bispo and B. Vilela helped with the map. IBAMA provided collecting permits (#14474) and the staff of the Serra da Bocaina National Park kindly provided logistical support and housing. This paper is part of DBP master’s thesis at the Biologia Animal graduate program, supported by FAPESP-São Paulo Research Foundation (#2008/55744-6) and CAPES-Coordenação de Aperfeiçoamento de Pessoal de nível Superior fellowships. We greatly acknowledge the input received from L. C. Schiesari and L. Casatti as members of DBP’s thesis examining committee. MVG and TG-S were supported by FAPESP doctoral fellowships (#2008/50575-1, 2008/58979-4). IAM was supported by a Jovem Pesquisador grant from FAPESP (01/13341-3 and 06/56007-0). During the final preparation of the manuscript DBP was supported by a CAPES-DS and CAPES-PDSE doctoral fellowships. We also thank all those who helped in the field and lab work. DCRF is a fellow of the Brazilian National Council for Scientific and Technological Development. Voucher specimens are housed at the tadpole collection of DZSJRP-UNESP.

